# Non-triazole agricultural fungicides indirectly select for triazole resistance in the human pathogen *Aspergillus fumigatus*

**DOI:** 10.64898/2026.05.28.728463

**Authors:** Hylke H. Kortenbosch, Spyros G. Kanellopoulos, Luisa Timmermanns, Francisca Reyes Marquez, Bas J. Zwaan, Ben Auxier, Eveline Snelders

## Abstract

The intensification of agriculture relies on chemical fungicides to manage crop disease^1,2^, leading to the evolution of resistance in plant pathogens.^3^ Fungicides have long half-lives, allowing them to remain active well beyond their intended targets and affect downstream ecosystems and agricultural practices. The saprophytic fungus *Aspergillus fumigatus* is an airborne ubiquitous fungus and an important human pathogen causing severe life-threatening invasive fungal disease.^4^ Selection pressure from agricultural triazoles, demethylase inhibitors (DMIs), has led to cross-resistance to clinical triazoles, as they share the same target gene, *cyp*51A.^5,6^ In the Netherlands^7,8^, most triazole resistance arises from two *cyp*51A haplotypes, the TR_34_ and TR_46_.^9,10^ Genomic surveys of *A. fumigatus* have shown that these triazole-resistance alleles often co-occur with resistance alleles to non-DMI classes, these include some of the dominant fungicide classes used in Europe such as quinone outside inhibitors (QoIs), and succinate dehydrogenase inhibitors (SDHIs).^11^ Hypothesizing that agricultural environments with non-DMI fungicides can indirectly select for triazole resistance, we used grass mesocosms to compete *A. fumigatus* isolates. We found that already at low concentrations, commonly found in agricultural residues, DMI, SDHI, and QoI fungicides can each independently increase the proportions of triazole resistant alleles. We show that the resistance alleles for each class are not intrinsically cross-resistant, indicating that their co-occurrences in allelic combinations produce this multi-resistance selection. Consistent with our mesocosms results, environmental samples contained high phenotypic (>20%) triazole resistance in heaps with only non-DMI fungicides. This work provides the first experimental and field evidence of selection for triazole resistance by non-triazole fungicides via genomic hitch-hiking. We thus predict that when novel fungicides are used in the same selective environment, novel resistance alleles will most likely be selected in isolates that have already accumulated resistance alleles to other fungicide classes. Because of linked resistance alleles, tackling selection and spread of environmental triazole resistance will require consideration of all fungicide classes.

**Graphical abstract:** 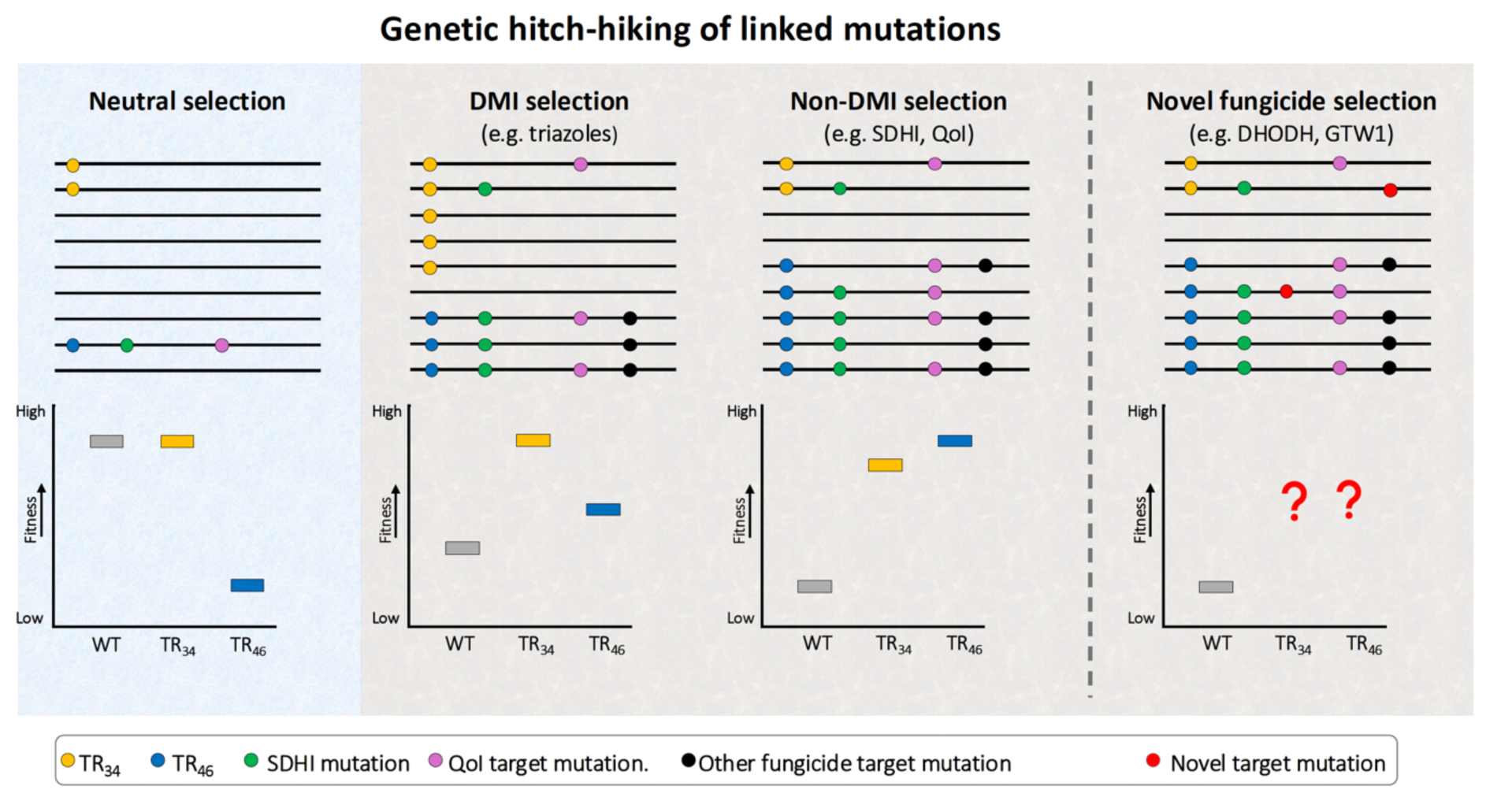

**Highlights:**

- Triazole-resistance alleles often co-occur with resistance alleles to non-DMI fungicide classes such as SDHI and QoI.
- DMI, SDHI, and QoI all independently increase the proportions of triazole resistant alleles at low concentrations (0.1 mg/kg) commonly found in agricultural residues.
- The DMI resistance allele TR_46_ has a high fitness only under fungicide selective conditions, whereas TR_34_ maintains its fitness also under fungicide free conditions.
- Beneficial alleles for one fungicide class increase in frequency and ‘hitch-hike’ with resistance alleles to other fungicide classes.
- Resistance alleles to novel fungicide classes will more likely be selected in isolates that already accumulated other fungicide resistance alleles.

## Results

First, to exclude the possibility that fungicide resistance alleles mechanistically cause cross resistance beyond their class and target genes, a mutant analysis was performed. To understand the selection of triazole resistance by multiple fungicide classes, we developed a three-dimensional grass substrate mesocosm to mimic the physical and nutrient conditions experienced by *A. fumigatus* in plant waste heaps (Figure S1). We constructed a pool of 187 *A. fumigatus* isolates originating from a recent Dutch nationwide air sampling study.^7^ The pool consisted of three groups of isolates with different haplotypes of the *cyp*51A promoter region: 50 triazole-resistant isolates carrying TR_34_, 50 triazole-resistant TR_46_, and 87 wild type isolates. Previous work showed that Dutch multi-triazole-resistant TR_34_/TR_46_ *cyp*51A haplotypes are associated with resistance to MBC, SDHI, and QoI classes, strongly suggesting an exposure history to multiple agricultural fungicides in these environmental hotspots.^15^ In addition to the *cyp*51A promoter region, no genomic variation was however considered for the selection of isolates, neither point mutations in *cyp*51A nor other fungal resistance alleles. We competed the pool of isolates in bulk across a concentration gradient of three fungicides: the DMI tebuconazole, the SDHI boscalid, and the QoI azoxystrobin.

### Single resistance alleles do not cause multi fungicide resistance

Mutant isolates were constructed in a wild type background with the following mutations; a TR_34_/L98H in the *cyp*51A gene and a H270Y in the *sdh*B gene. The G143A allele in the *cyt*B gene was not constructed as it is located in the mitochondrial genome and no methodological approached are available yet. All CRISPRcas9 constructed mutants with their single resistance alleles were phenotypically tested for antifungal resistance to the following classes of fungicides; DMI, SDHI, and QoI (Figure S2). With breakpoints lacking for fungicides in *A. fumigatus*, resistance was determined based on growth at concentrations that exceeded the threshold assumed to distinguish susceptible from resistant populations. Resistance was observed only when the specific target resistance allele was present, and none of the mutant isolates exhibited cross-resistance to other fungicide classes (Figure S2). Therefore, the selection of triazole resistant *cyp*51A alleles cannot be the result of single resistance alleles that are cross-resistant beyond their own class of fungicides.

### Selection of *cyp*51A and *sdh*B resistance alleles in grass mesocosm by fungicides

Consistent with the predicted DMI resistance of the TR_34_ and TR_46_ alleles, tebuconazole strongly selected both *cyp*51A resistance haplotypes relative to fungicide-free conditions in the grass mesocosm (Fig. 1A, top panel). In particular, the TR_46_ haplotype had significantly lower fitness (β = −1.16, SE = 0.141, p = < 0.01) under fungicide free and increasing fungicide-conditions. Even at 10 mg/kg of tebuconazole, TR_46_ fitness did not exceed that of wild type alleles (β = − 0.00145, SE = 0.141, p = 1.00). Interestingly, in contrast to tebuconazole, boscalid and azoxystrobin treatment produced similar results as tebuconazole, with TR_34_ haplotypes having slightly higher fitness across all fungicide concentrations, but here TR_46_ transitioned from low to high fitness at concentrations above 0.1 mg/kg (Fig. 1A, middle and bottom panel). At the highest concentrations, the TR_46_ isolates had a higher fitness than TR_34_ isolates, although this was statistically significantly different only for azoxystrobin. To understand the multi fungicide resistance nature of selection, we also measured fitness by the frequency of the H270Y resistance allele in the *sdh*B gene, which provides resistance to the SDHI boscalid (Fig.1B). In contrast to *cyp*51A alleles, we did not preselect isolates based on their *sdh*B genotype.

**Figure 1.**
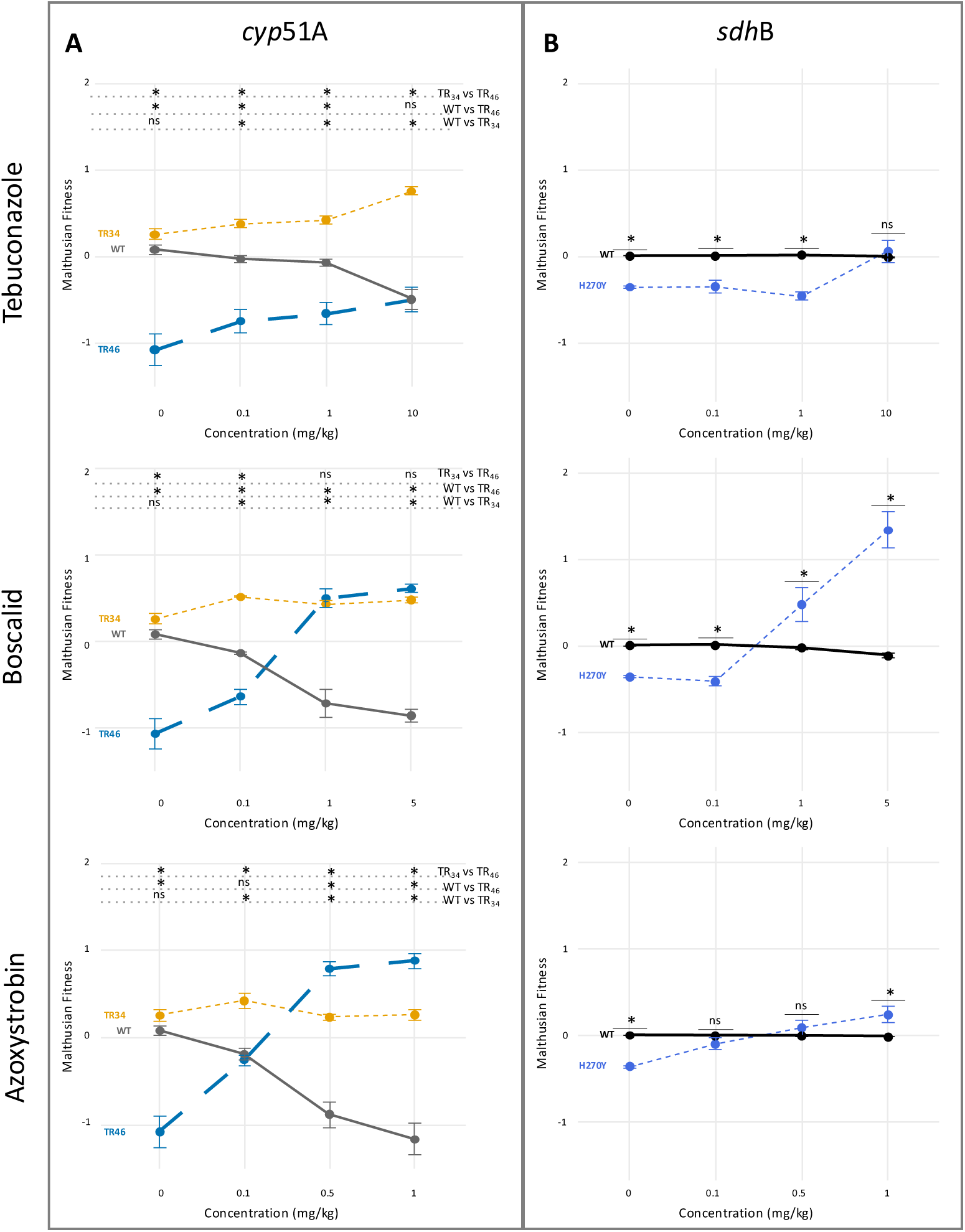
The fitness of *A. fumigatus* in grass mesocosms shows resistance allele specific rather than fungicide specific dynamics. A) The fitness of the *cyp*51A haplotypes measured by allele frequency across increasing concentrations of tebuconazole (top), boscalid (middle) and azoxystrobin (bottom) (P1) relative to the allele frequency of the starting suspension (P0). Tebuconazole selects for both TR alleles, but TR46 shows a pronounced fitness cost under conditions free of fungicides and low-concentration conditions. In contrast, under boscalid and azoxystrobin, TR46 changes from low to high fitness at ≥0.1 mg/kg, even exceeding TR34 at the highest concentrations (p < 0.01. B) The fitness of sdhB alleles at increasing concentrations of fungicides. The H270Y allele is strongly selected only under boscalid (≥0.1 mg/kg), while selection under tebuconazole and azoxystrobin is weak or absent except at the highest concentration of azoxystrobin. Similar to TR46, H270Y carries a fitness cost in fungicide-free conditions. *Post hoc* pairwise comparisons were performed using *t*-tests on estimated marginal means with Bonferroni correction for multiple testing. (* = p < 0.05, ns = not significant). The pairwise differences between the *cyp*51A alleles can be found in Table S2, and for the H270Y *sdh*B allele in Table S3. The raw proportions underlying the fitness data are shown in Figure S3.

Consequently, 3.6% of the reads in our starting inoculum carried the H270Y variant (Fig. S3). Under boscalid treatment, which directly selects for *sdh*B resistance, the resistant allele was strongly selected for from the pool of isolates at concentrations above 0.1 mg/kg (Fig. 1B, middle panel). Under tebuconazole and azoxystrobin treatment the selection was weaker, with a significant difference to the wild type only found at the highest (1 mg/kg) concentration of azoxystrobin, while the highest concentration of tebuconazole was only sufficient to raise the fitness to be equal with the wild type allele (Fig 1B, top and bottom panel). Similar to the TR_46_ *cyp*51A allele, the H270Y *sdh*B allele was associated with a significant fitness cost under fungicide-free conditions (Fig. 1B). Fitness for the G143A *cytB* variant was not measured as it is located in the mitochondrial genome, and the ratio between nuclear and mitochondrial genome is not a fixed parameter. Fitness measurements in these grass mesocosms compared to traditional flat solid agar showed large differences (Fig. S4). For *cyp*51A, the fitness of the wild type under high-fungicide conditions for all three fungicides on agar was below -3, translating to a ∼95% proportional decrease in a single generation, far lower than in our grass mesocosm. All pairwise differences between fitness on agar *versus* grass can be found in Table S4.

### Selection of individual triazole resistant *cyp*51A alleles by fungicides

Because our haplotypes are represented by pools of isolates, the fitness differences observed could be driven either by a few resistant individuals with high fitness in our conditions or by consistent patterns of the subpopulations. To test this, we used existing segregating markers in the *cyp*51A gene that allowed us to decompose the population into 32 discrete haplotypes; 18 wild type, eight TR_34_, and six TR_46_. Comparing the wild type and TR_34_ haplotypes (Fig. 2A) shows that five of eight TR_34_ haplotypes have a fitness advantage over all wild type alleles above 0.5 mg/kg azoxystrobin and for boscalid, the same applies to seven of eight TR_34_ haplotypes from 1 mg/kg and above. All individual alleles, wild type or TR_34_, show the same upward or downward trends in increasing concentrations of fungicides and therefore show consistent results. On tebuconazole the separation in fitness between wild type and TR_34_ is less clear, although there is a clear divergent trend between the two groups beyond 1 mg/kg. Comparing the wild type and TR_46_ haplotypes (Fig. 2B), five out of six TR_46_ haplotypes outcompete all wild type haplotypes at both 0.5 mg/kg azoxystrobin and 1 mg/kg boscalid but this is not observed for tebuconazole (up to 1 mg/kg). Comparing the TR_34_ and TR_46_ haplotypes (Fig. 2C) shows that the TR_46_ haplotypes generally have a lower fitness in the absence of fungicides or on tebuconazole but have similar fitness at higher concentrations of azoxystrobin and boscalid. To indicate the general correlation between different haplotypes, the fitness of all identifiable *cyp*51A haplotypes at the highest rates of azoxystrobin (1 mg/kg) *versus* boscalid (5 mg/kg) was strongly correlated (R^2^ = 0.84), fully separating the wild type *cyp*51A haplotypes from the TR haplotypes (Figure S5).

**Figure 2.**
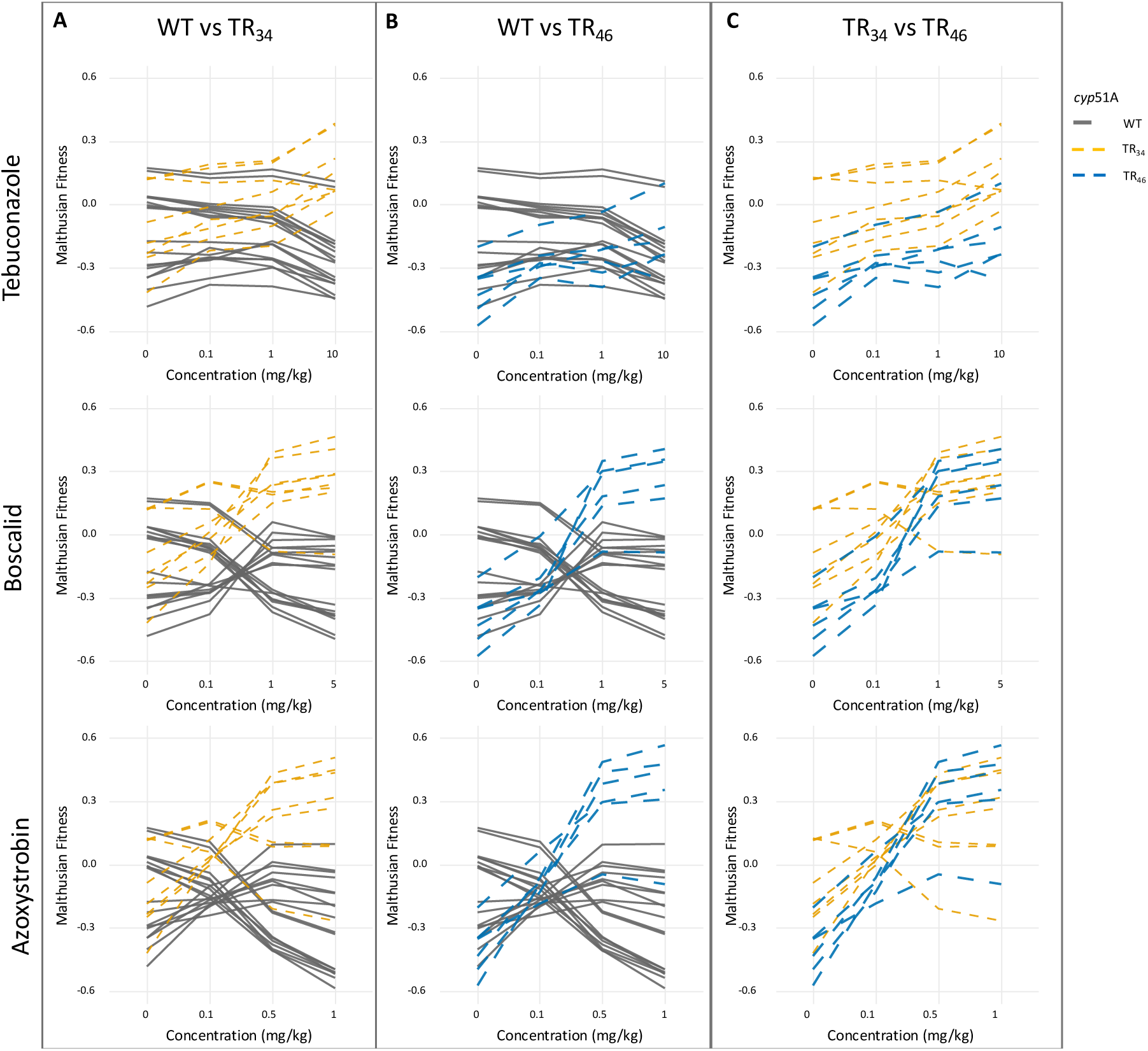
Consistent multi-fungicide selection across individual triazole resistant *cyp*51A alleles. Fitness of distinct *cyp*51A sequence haplotypes in increasing concentrations of tebuconazole, boscalid, and azoxystrobin in the grass mesocosm (Wild Type (WT) (n=18) indicated by a solid grey line, TR34 (n=6) by a dashed yellow line, TR46 (n=8) by a dashed blue line). A) TR34 haplotypes largely outperform wild type haplotypes at higher fungicide concentrations, and most TR34 variants show a fitness advantage above 0.5 mg/kg azoxystrobin and ≥1 mg/kg of boscalid and tebuconazole. B) TR46 haplotypes outperform wild type at azoxystrobin and boscalid at ≥0.5–1 mg/kg, but not under tebuconazole, where no consistent advantage is observed. C) Direct comparison of TR34 and TR46 haplotypes shows that TR46 generally has lower fitness in fungicide-free conditions but reaches comparable fitness to TR34 at higher azoxystrobin and boscalid concentrations, while remaining less competitive under tebuconazole.

### Multi-fungicide detection in field samples with triazole resistance

To validate the results of grass mesocosms and testing the hypothesis of multi fungicide selection in natural settings, we paired triazole-resistance and HPLC fungicide data from a previous surveillance study (Supplementary material).^12^ Here, we investigate whether fungicide concentrations that select for triazole resistance in our grass mesocosms also co-occur with high phenotypic resistance fractions in the environment, particularly when DMI fungicides are not present. Eighteen of these samples had sufficient density of *A. fumigatus* (≥5,000 CFU/gr) to indicate growth in the substrate (Supplemental Material), with phenotypic triazole resistance fractions between zero and 60% (mean fraction 21,6%). Of the seven high-triazole resistance samples (>20% resistance), only three had notable DMI residues that can explain selection of resistance. (Figure 3A). Three other samples with phenotypic high triazole resistance showed selective QoI levels (0,108; 0,015; and 0,08 mg/kg Figure 3B), and one with selective methyl benzimidazole carbamate (MBC) levels (0,058 mg/kg) (Figure 3C). Five of the high triazole resistance samples and two of the low-resistance piles had substantial SDHIs selective levels that ranged between 0,042 and 5,61 mg/kg. The combination all four fungicide classes better explained the selection of seven of the eighteen positive triazole resistant field samples than assessing DMI levels only (Figure 3E).

**Figure 3.**
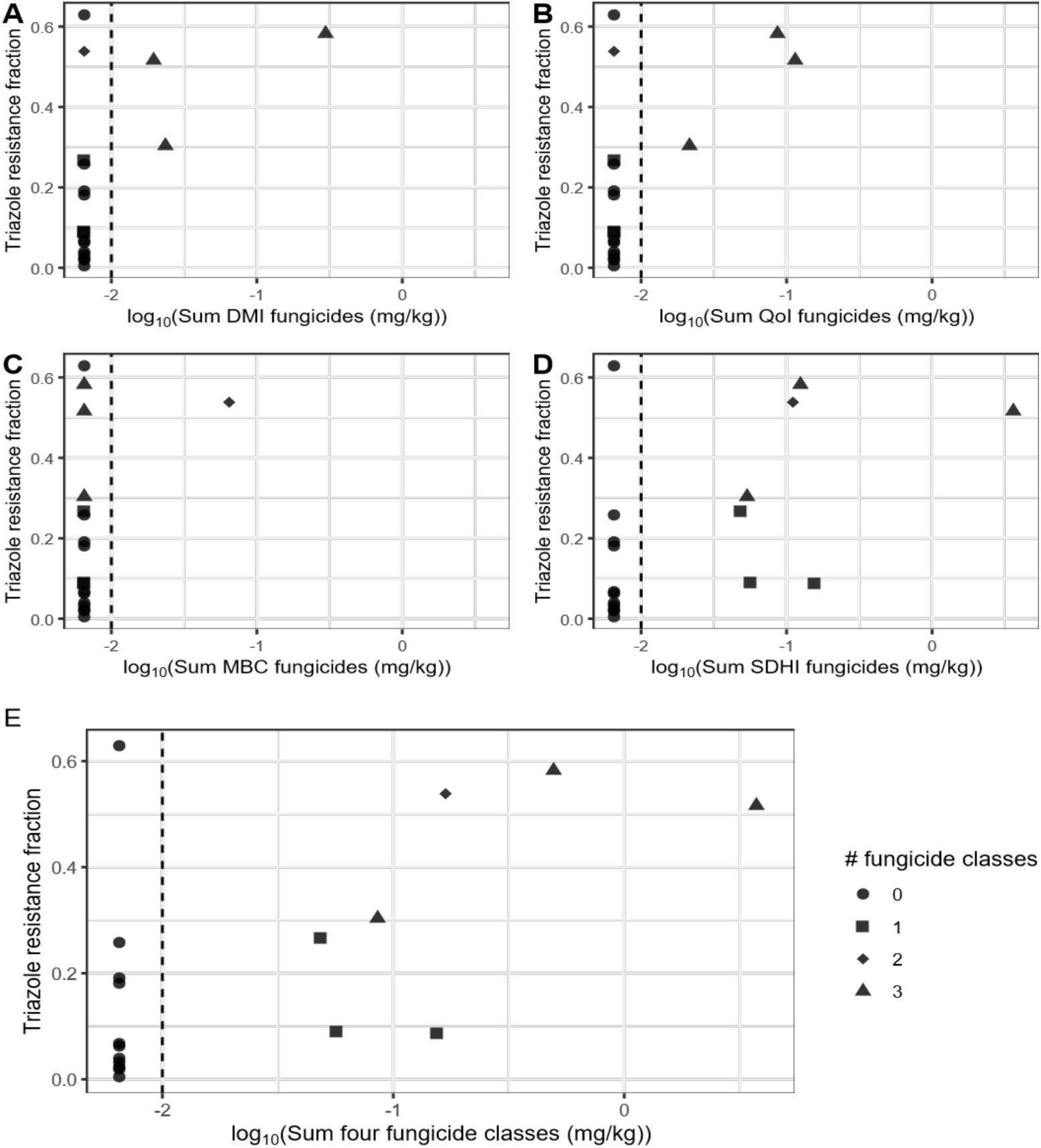
Multi fungicide detection in field samples with phenotypic resistance to triazoles. Comparisons of the concentrations of four fungicide classes in 18 environmental samples against the mean *A. fumigatus* triazole resistance fraction of these samples (Supplemental material). The dashed vertical line marks the detection limit (0.01 mg/kg) of the fungicide measurements. The fungicide classes shown in the subfigures are A) DMIs, which include triazoles and only show for three samples a correlation with presence of DMI fungicides and triazole resistance detection. High triazole resistance fractions (>0.2) can also be detected in the presence of non-DMI class fungicides with in; B) QoIs, which include boscalid, C) MBCs, and D) SDHIs, which include azoxystrobin. And finally in E) the sum (mg/kg) of the total concentration of fungicides detected with the shapes representing the number of fungicide classes present in total in each sample. Here, all high triazole resistance fractions, except for one sample, co-occur with the detection of one or more fungicide classes.

## Discussion

*A. fumigatus* isolates often carry resistance alleles to several fungicide classes to which this ubiquitous fungus is exposed. In recent years, the TR_34_ and TR_46_ haplotypes have been found to carry resistance alleles more frequently to non-DMI classes than triazole-sensitive isolates.^13–15^ A recent study that included 729 *A. fumigatus* isolates predominantly from the United States found that the H270Y variant of the *sdh*B gene occurred exclusively in TR_34_/TR_46_ isolates at a frequency of ∼15% for TR_34_ and ∼44% for TR_46_ isolates.^13^ Additionally, many of the H270Y-carrying isolates, carried the G143A allele of *cyt*B, which confers resistance to several QoIs. In an independent pool of Dutch clinical and environmental TR_34_/TR_46_ isolates phenotypic evidence was shown for multi-fungicide resistance to both MBC and QoI-class fungicides.^15^ Even though we did not construct the pool of isolates in this study based on resistance alleles other than the *cyp*51A TR alleles and thereby DMI resistance, our grass mesocosms results are consistent with selection of preexisting DMI resistance alleles by the QoI fungicide azoxystrobin and the SDHI boscalid. This work shows that exposure to different agricultural fungicide classes leads to selection for the DMI resistance alleles TR_34_ and TR_46_. More specifically, fitness of *A. fumigatus* resistance alleles in fungicide exposed grass mesocosms show allele specific rather than fungicide-class-specific dynamics.

Our work provides concrete evidence in current *A. fumigatus* populations that selection by genomic association of resistance alleles, also known as genetic hitch-hiking, can explain how the application of one fungicide can select resistance alleles to fungicides with different modes of action. We show that within an individual, the *cyp*51A or *sdh*B resistance alleles do not directly produce cross-resistance beyond DMI or SDHI-class fungicides, respectively. However, at a population level, we see similar fitness responses to boscalid and azoxystrobin by *cyp*51A resistance alleles, indicating strong correlation of these alleles in multi-fungicide resistance phenotypes. Although often interpreted as the physical linkage of genes on the same chromosome, genetic linkage is simply the statistical co-occurrence of alleles.^16^ Our results demand a reassessment of what conditions select for triazole resistance. Studies that define triazole resistance ‘hotspots’ have logically focused on the presence of triazoles and other DMI-class fungicides as a necessary condition.^17–19^ Our grass mesocosms and observational field data confirm that non-triazole fungicides select for triazole resistance within the existing genetic variation of *A. fumigatus* under environmentally relevant growth conditions. We provide empirical evidence that, at concentrations detected in agricultural waste, the QoI fungicide azoxystrobin and the SDHI fungicide boscalid each select for triazole resistance in *A. fumigatus*. Because we observed selection at the lowest concentration used, we cannot yet define the minimal selective concentrations for these compounds. Similarly, we have not explored the consequences of mixtures of multiple fungicide classes; whether the selective effects can be explained by a simple additive model or whether synergistic effects increase the selective pressure. In 2025, EU agencies produced a report requested by the European Commission reviewing current evidence and providing conclusions and recommendations on the impact of azole fungicide us and the development of resistance in *A. fumigatus*.^20^ It was concluded that ‘hotspots’ for azole resistance selection are those with substrates supporting the growth of *A. fumigatus* and that these must include levels of DMIs exceeding the PNEC_res_ (Predicted No Effect Concentration for resistance selection) value. We now strongly advocate including other classes of fungicides and expanding the range of exposure concentrations and combinations when developing risk assessments or performing *in vitro* experimental work with a necessary broader focus than only DMIs.

Although both TR_34_ and TR_46_ provide resistance to triazole, we found striking differences in fitness between these groups in the absence of fungicides. TR_46_ is the dominant triazole resistance haplotype in Dutch multi-fungicide flower bulb waste heaps^18^, yet our recent air sampling campaign showed that it was rare outside of these environments.^7^ Across 355 sites TR_34_ had a uniform but low (∼5%) occurrence while TR_46_ showed high local peaks (up to ∼20%) that correlated with specific agricultural land uses.^9^ Because TR_46_ isolates are more likely to be multi-fungicide resistant^15,21^, we can now explain that the competitive advantage of TR_46_ isolates is limited to multi-fungicide environments, while in the absence of fungicides the TR_34_ isolates are better competitors. These fitness differences are consistent with recent pairwise agar-based competition experiments where environmental TR_46_ isolates had lower fitness in the absence of fungicides compared to both wild type and TR_34_ isolates of environmental origin.^22^ Here, Chen *et al.* showed, by using genetic transformation, that the fitness costs in the TR_46_ triazole-resistant isolates was not caused by the *cyp*51A alleles themselves, implying the presence of deleterious alleles elsewhere in the genome.^22^ However, a study on the global *A. fumigatus* population did not find a fitness cost for TR_46_ isolates when competed in liquid growth media.^23^

Our experimental results have clear policy implications. If we maintain current fungicide usage patterns, the existing multi-resistant *A. fumigatus* strains are in the ideal ecological position to continue accumulating resistance mechanisms to novel modes of action, with the TR_46_ allele as most likely to accumulate these novel alleles.^24^ The accumulation of novel resistance mechanisms is of particular concern for the novel DHODH (dihydroorotate dehydrogenase) and GWT1 (Glycosylphosphatidylinositol-anchor biosynthesis protein 1) antifungal classes. Both of these classes are expected to see widespread dual clinical and agricultural use alongside existing antifungals.^25,26^ Consequently, strains with novel resistance alleles will most likely also carry triazole resistance mechanisms, complicating treatment through frequent multi-fungicide resistance. Non-DMI fungicides can select for triazole *cyp*51A resistance alleles, and this new evidence needs to inform policymakers in developing and evaluating effective measures to mitigate environmental selection for triazole resistance. While in the short term the focus may be on waste management strategies or reduction of TR_46_ populations, in the long term an overall reduction of fungicide use, not just DMIs or dual use compounds, is needed to reduce selection for and the burden of triazole resistance in *Aspergillus fumigatus*.

## Data and code availability

Raw data and analysis code used in this study, and required to re-analyse the data, will be made available.

## Acknowledgements

The authors thank Bo Briggeman for facilitating the use of the ‘Schimmelradar’ isolates and Merijn Pieterse for troubleshooting the molecular work with the grass substrate, especially the DNA extraction protocol. We thank Peter Leendertse for providing us with broader fungicide data beyond the triazole data reported in Leendertse, Gommer & van Beek, 2021. Finally, we would like to thank Dr. Michael Bottery for providing critical feedback on our manuscript. Primary research funding was provided through the Netherlands Organisation for Scientific Research (NWO) under the Groen III program (GROEN.2019.002). FRM was supported by an Aspasia project (015.015.028). The authors declare no competing interests.

## Author contributions

Conceptualisation: HK, SK, BA, and ES; Data curation: HK, SK, and BA; Formal analysis: HK, SK, and BA; Funding acquisition: ES, BZ; Investigation: HK, LT, and FRM; Methodology: HK, SK, LT, FRM, and BA; Project administration: HK and ES; Software: HK, SK, and BA; Supervision: BA, and ES; Visualisation: HK; Writing-original draft: ES; BA; HK Writing-review and editing: ES, BA, HK, SK, FRM, LT and BZ.

## Methods

### Protoplast preparation

Our CRISPR-Cas9 transformation process was based upon Abdallah *et. al.* 2017.^25^ We used *A. fumigatus* transformation recipient strains Afir974 and Afir964.^26^ Spores of either strain (2x10⁸) were inoculated into 40 mL of complete media (CM)^27^ and incubated at 30 °C and 100 rpm shaking for 16 hours. After incubation, the cultures were filtered through Miracloth (Merck) and washed twice with 10 mL CM. The mycelium was then transferred to a CM-protoplasting solution (5% VinoTaste (Novo Nordisk) 16 mL CM, 16 mL of 1M KCl-0.1M Citric acid solution). The flask was incubated at 30°C and 100 rpm for 3 hours. The solution was then filtered through a 40 µm cell strainer and centrifuged at 1,800 g for 10 minutes. The supernatant was discarded, and the protoplasts were resuspended in 2 mL of 0.6 M KCl. The protoplasts were centrifuged again at 2,400 g for 3 minutes. The supernatant was removed, and the pellet was washed twice with 0.6 M KCl. Finally, the protoplasts were resuspended in 1 mL of final solution (0.6 M KCl; 50 mM CaCl₂) and adjusted to a final concentration of 5 × 10⁶ protoplasts/mL.

### GuideRNA and Single-Stranded Donor DNA design

For the transformation, guideRNAs (gRNA) were designed using software, EuPaGDT (Supplementary Table S1).^28^ The crisprRNAs (crRNA) with the highest QC scores and 5’ orientation was selected and ordered (Alt-R^TM^ CRISPR-cas9, Integrated DNA Technologies (IDT)). As donor DNA, we used ∼100bp single stranded donor DNA (IDT) centered on the desired genetic change with 50 bp upstream and 50 bp downstream flanking the target region (Supplementary Table S1). RiboNucleorotein Particle (RNP) complexes were assembled^25^ and for generation of the gRNA, each crRNA was hybridized with the trans-activating crisprRNA (tracrRNA) by combining them in a 1:1 ratio in nuclease-free duplex buffer (IDT) to a final concentration of 33 μM. The mixture was brought to 95 °C and cooled down to 25 °C at a rate of 0.1 °C/s. For the formation of the RNP complex 1.5 μL of the 33 μM duplex solution, 1.5 μL of Cas9 nuclease (1 μg/μL, IDT), 23.5 μL of Cas9 working buffer (20 mM HEPES, 150 mM KCl, pH 7.5) were combined to a final volume of 26.5 μL RNP reaction mixture.

### Transformation

A standard transformation protocol was followed for the transformation of *A. fumigatus*.^25^ Initially, 200 µl of protoplasts were transferred to a 1.5 mL tube containing the 26.5 μL of RNP reaction mixture. Additionally, 1 μg of the pMA171 plasmid^29^ was added and 25 µl of polyethylene glycol (PEG)-CaCl_2_ buffer (60% [wt/vol] PEG 3350, 50 mM CaCl_2_⋅H_2_O, 450 mM Tris-HCl, pH 7.5) incubating on ice for 30 min. Next, 1.25 mL PEG-CaCl_2_ were added followed by a 20-minute incubation period at room temperature. Transformed protoplasts were spread across eight YPS agar plates (yeast extract 20 g/L, peptone 5 g/L, sucrose 1.0 M, Tris base 5 mM, agar 15 g/L, pH 6.0) supplemented with 250 mg/L hygromycin B (Duchefa Biochemie, The Netherlands) and incubated for 72 hours at 37°C. To confirm genomic DNA editing, the colonies that grew on YPS-hygromycin plates, were transferred to a 96 deep-well plates with CM and after incubation for 48 hours at 37 °C spores were obtained. PCR-Quality DNA isolation was performed for each sample ^30^ and the sequence validation of the transformants was done by sanger sequencing (Mix2Seq, Eurofins) the PCR products obtained using primers flanking the site of genomic edition. For *sdh*B, primers sdhB_Check_Fw 5’ TTGATCCCTTTCGGCTCGTC 3’ and sdhB_Check_Rv 5’ TGTCCAATTTCCCTGTGCGT 3’ were used. For *cyp*51A, one set of primers was used to check the TR insertion Fw (pA7) 5’ TCATATGTTGCTCAGCGG 3’ and Rv (pA5) 5’ TCTCTGCACGCAAAGAAGAAC 3’.^31^ A second set was used to check for the L98H mutation; Fw (365.1) 5’ ATGGTGCCGATGCTATGG 3’ and Rv (1.77) 5’ CTGCACATACAAAAACGGCCAGCAA 3’.^32^

### Antifungal susceptibility testing of CRISPRcas9 mutant isolates

Successful CRISPRcas9 mutant isolates, wild type (AF293) and the transformation recipient isolates (Afir974 and Afir 964) were assessed for antifungal susceptibility. Five μL of a 10⁷ spores/mL suspension of each strain was spotted onto 9 cm plates with minimal medium (MM)^27^ containing either no fungicide or supplemented with 4 mg/L tebuconazole, 5 mg/L boscalid, or 5 mg/L azoxystrobin and incubated for 92 h at 37°C.

### Isolate pool preparation

To prepare the inoculum, we first made individual highly concentrated suspensions of the asexual conidiospores for all isolates in PBS-0.05% (v/v) Tween-80 (pH 7.4). We then combined 100 µL of each isolate suspension belonging to the same variant group: wild type, TR_34_, and TR_46_. We determined the spore concentration of each of the three pools using a CASY Cell Counter & Analyzer (OMNI Life Science, Bremen, Germany). Based on the spore concentrations, we then calculated the volumes of undiluted suspensions required to make 50 mL of inoculum with 500.000 conidia/mL, containing 46% wild type, 27% TR_34_ and 27% TR_46_. We chose this ratio to approximately equally represent every isolate in the inoculum. Of the resulting ∼500 µL of concentrated inoculum, ∼334 µL was used to dilute to 500.000 spores/mL. The remaining ∼167 µL of concentrated suspension was used to determine the starting inoculum (P_0_).

### Grass substrate and fungicides

As the primary growth substrate, we used freeze-dried grass cuttings from the lawn of a residential backyard in Wageningen (Coordinates: 51.95610°N, 5.64394°E (WGS84)). We confirmed that the grass was free of fungicides, including triazoles, QoIs and SDHIs, down to 0.01 µg/kg via High-Performance Liquid Chromatography (GC-MS/MS and LC MS/MS, Eurofins lab Zeeuws Vlaanderen, The Netherlands). The grass was frozen at -20 °C within 24 hours of collection for later processing. Three months ahead of the competition experiment, the grass was freeze-dried and subsequently sieved down through a 1 cm grinder filter. The freeze-dried grass was then stored at room temperature in a sealed container for later use. The competition experiments were performed in 30 mL glass bottles with metal caps and silicon seals. At this stage, we prepared fungicide solutions by dissolving the fungicides in demineralised water from dimethyl sulfoxide (DMSO) (Merck) stocks. We added 4 mL of these solutions to their respective grass bottles to both rehydrate the grass and add the fungicides at the desired concentrations. For the agar treatments, we used 10 mL of Malt extract agar (MEA) (Sigma-Aldrich) per bottle (1.5% (w/v) agar), supplemented with 1 mg/L CuSO_4_ (Sigma-Aldrich). For the triazole tebuconazole (Sigma-Aldrich), the final concentrations were 0.1 mg/kg, 1 mg/kg, and 10 mg/kg; for the QoI azoxystrobin (Sigma-Aldrich), 0.1 mg/kg, 0.5 mg/kg, and 1 mg/kg; and for the SDHI boscalid (Riedel de Haën), 0.1 mg/kg, 1 mg/kg, and 5 mg/kg. We selected the fungicide concentrations based on their relevance, the fungicide-specific solubilities in water, and the reduction of wild type *cyp*51A observed in earlier pilot experiments. For the true negative and growth control treatments, 4 mL of demineralised water was added to rehydrate the material. The bottles were subsequently autoclaved to sterilise the material and disperse the fungicides throughout the material. For the agar treatments, we added the highest concentration of each of the three fungicides in grass to each corresponding set of agar bottles. Due to the autoclaving step needed to spread the fungicides through the grass substrate, representative fungicides were selected both for high thermal stability (> 121 °C) and their different modes of action. Additionally, it was confirmed that all three compounds retain their inhibitory effect on sensitive isolates after autoclaving in MEA medium (data not shown).

### Spore inoculation, incubation and harvest

Samples were inoculated with 100 µL of mixed inoculum (containing ∼50.000 spores). To spread the spores through the grass substrate, the suspension was added by gently dribbling droplets of suspension into the bottle while moving the pipette tip in a circular motion. For the agar samples, the bottles were also briefly swirled to spread the liquid across the surface. *A. fumigatus* can grow in the grass substrate with the bottles fully closed but will grow slowly and show minimal sporulation. We therefore screwed the bottle caps on loosely by only half a turn to allow for gas exchange. Bottles were distributed across two boxes with their lids 2 cm open to facilitate gas exchange. Samples 1 to 34 were placed in one box, and samples 35 to 68 in the other box. All samples were incubated at 37 °C for 4 days, at which stage there was abundant, visible spore production on the grass. To harvest the spores from the bottles, 15 mL of PBS-0.05% (v/v) Tween-80 (pH 7.4) was added to all bottles. The bottles were subsequently shaken from side to side with the bottle caps firmly closed at a frequency of 250 times per minute for 5 minutes on a GFL 3018 shaker (Gemini BV) to suspend the spores. Immediately before collecting 1.5 mL of spore suspension from the bottles, each bottle was again vortexed briefly. In case the hydrophobic mycelium caused the grass to float, the spores were left to settle for a few minutes to prevent cross-contamination with airborne spores in the biosafety cabinet. Then, the grass was pushed down into the liquid and the bottle vortexed again. Spores were otherwise collected immediately after vortexing each sample. The harvested suspensions were moved into 1.5 mL tubes and centrifuged for 3 min at 5000 rpm. To ensure sufficient spores for DNA extraction, the aim here was to have firm (∼3 mm) pellets at the bottom of all Eppendorf tubes. If no pellet was visible at the bottom of the tube, the supernatant was removed, and 1.5 mL of spore suspension from the affected sample was added again on top.

### Molecular determination of alleles

Before and after four days of growth at 37°C, we used *cyp*51A and *sdh*B genotypes as markers for triazole resistance and SDHI resistance, respectively, and tracked proportional shifts using nanopore sequencing. For the DNA extraction of the P_0_ spore suspension, we used a heat shock protocol.^33^ The heat shock protocol works well on clean *A. fumigatus* spore suspensions but proved ineffective to extract and subsequently amplify DNA from the grass substrate, possibly due to organic inhibitors released from the grass. Therefore, we used a modified version of an established DNA extraction protocol using ‘breaking’ buffer, which uses phenol-chloroform to remove metabolites from the sample.^34^ At the start of the protocol, rather than transferring spores into the breaking buffer with a cotton swab, we spun down the harvested suspensions from the grass samples for 3 min at 5000 rpm. We then removed the supernatant and added the breaking buffer and sterile sand/beads on top of the pellet. DNA extraction was performed as previously described.^34^ To capture *cyp*51A variation, a ∼2200 bp fragment was amplified that includes the promoter and the coding region. We used nanopore compatible tailed primers to amplify the *cyp*51A gene: Fw 5’TTTCTGTTGGTGCTGATATTGCTCATATGTTGCTCAGCGG-3’ and Rv 5’-ACTTGCCTGTCGCTCTATCTTCCTGTCTCACTTGGATGTG-3’ (nanopore tails in bold). To capture the variation in the *sdh*B gene, we amplified a ∼750 bp fragment including the H270Y resistance SNP using Fw 5’-TTTCTGTTGGTGCTGATATTGCACACCAAGACCGAGGATGTG-3’ and Rv 5’-ACTTGCCTGTCGCTCTATCTTCTCAATTGCCCCAGAAGAAAACG-3’ primers, nanopore in bold. VeriFi Mix Red (Pcr Biosystems) was used for the PCR reactions. 25 µL PCR reactions were done according to the manufacturer’s instructions, with the addition of bovine serum albumin to a final concentration of 0.4 mg/mL and DMSO at 4% final concentration. For the P_0_ amplification, we used 2 µL of undiluted heat shock product as template. For the agar samples, we used 1 µL or 2 µL undiluted template DNA to amplify the *cyp*51A and *sdh*B fragments, respectively. For all grass samples, both PCR products were obtained using 3 µl of 10x diluted DNA per reaction as template. The PCR programme for the *cyp*51A fragment was as follows 95°C – 1 min, 35 cycles of 95°C – 15 s, 60°C – 15 s, 72°C – 1 min, and one cycle of 72°C – 5 min. After finalising the cycles, the temperature was brought down to 12°C. The PCR programme for the *sdh*B fragment was 95°C – 3 min, 30 cycles of 95°C – 30 s, 65°C – 30 s, 72°C – 2:30 min, and one cycle of 72°C – 5 min. All PCR reactions were checked for successful amplification by visualisation using gel electrophoresis. The true negative (no-growth) samples showed no bands, and at this stage were omitted from further analyses. To mitigate the effect of PCR amplification bias on our samples, each PCR reaction was performed three times using different thermocyclers and pooled, per fragment per sample. For the PCR fragments amplified from agar samples 12 µL per PCR reaction were pooled, to 36 µL total; all other samples were pooled with 8 µL per reaction to a total of 24 µL. The starting inoculum subsamples were vital for fitness estimations. We therefore pooled the three starting inoculum reactions in duplicate for each fragment for both subsamples. At this point, we placed the samples in a 96-well format, ordered by the random number they had been assigned at the sample incubation step. The samples were subsequently purified using Sera-Mag SpeedBeads (Cytiva 45152105050250) resuspended in 15 µl volume and quantified using Quant-iT PicoGreen (Invitrogen). Because Nanopore sequencing favours shorter fragments, we pooled the fragments at a molar ratio of 0.75:1 *sdh*B:*cyp*51A for each of the samples before barcoding them using the PCR Barcoding Expansion 1-96 (EXP-PBC096 from Oxford Nanopore Technologies). The barcoding, adaptor ligation using the kit V14 (SQK-LSK114), and library preparation followed the protocols available from Oxford Nanopore Technology. DNA concentrations in all these steps, except for the final pooled library, were measured by Invitrogen Quant-iT PicoGreen. For DNA quantification of the final pooled library, we used the Qubit 2.0 fluorometer using the dsDNA High Sensitivity Assay kit (Thermo Fisher).

The raw sequence data were aligned to the *A. fumigatus* reference genome of Af293 (GCA 000002655.1) with minimap2.^35^ Reads aligned to *cyp*51A were genotyped using pysam^36,37^ for the presence of the known alleles L98H, Y121F, T289A, as well as four other common synonymous SNPs. Reads aligning to *sdh*B (XM 748512.2) were genotyped for H270Y, as well as four other synonymous SNPs in the coding region. For both genes, common alleles were identified from a genomic analysis of global datasets containing ∼1200 genomes (unpublished data), selecting for alleles above 3% in the population. Reads were genotyped for each of these alleles, and the result was exported as a csv file. To avoid including sequencing errors, we produced a set of trusted haplotypes observed at least 10 times in both subsamples of the starting inoculum. From all samples, only reads that matched one of these trusted haplotypes were retained for subsequent analysis. Subsequently, the proportions among the reads of both the main *cyp*51A variant groups and the *sdh*B resistance variant were determined for each sample for the main results as presented in Figure 1. These proportions were then used to calculate Malthusian fitness through the change in the sample proportion relative to the proportions in the starting inoculum samples, assuming at least one generation had passed (see Equation 1).

### Field sampling fungicide resistance data

To evaluate environmental triazole resistance from a multi-fungicide perspective, we used data from an earlier surveillance study^12^ (Supplemental Material). In that study, samples were taken from plant waste heaps at various stages of decomposition with different waste materials, including green waste, wood chippings, flower bulb waste, and grass cuttings. Through selective culturing, the *A. fumigatus* density in colony-forming units (CFU/g) of these samples was determined.^38^ Concurrently, phenotypic triazole resistance fractions for itraconazole and tebuconazole were estimated for *A. fumigatus* in these samples by selectively culturing on plates supplemented with these triazoles. The fungicide concentrations for all fungicides permitted for use in the Netherlands were also measured in the samples, although only the total triazole concentration per sample was listed in the original report.^12^ These measurements were done using Gas Chromatography-tandem Mass Spectrometry (GC-MS/MS) and Liquid Chromatography-tandem Mass Spectrometry (LC-MS/MS) detection methods. The detection limit of the measurements was at 0.01 mg/kg. The detected fungicides were classified according to their respective classification on the Fungicide Resistance Action Committee (FRAC) Code List 2025.^39^ We included samples from this study in our visualization when the samples had a reported *A. fumigatus* density of at least 5,000 CFU/g, indicative of active *A. fumigatus* re-growth at the sampled site. Furthermore, to be confident that the *A. fumigatus* in the sample had been in contact with the measured fungicides during growth and reproduction, only samples from plant waste heaps that were over one day old were included.

### Statistical analyses and data visualisation

All statistical analyses were done in R (version 4.4.0).^40^ To model the fitness of both the *cyp*51A and *sdh*B variant groups under the different fungicide conditions, we used linear mixed-effects models (LMMs) of the lme4 package.^41^ The *cyp*51A alleles, split on their promoter region haplotype; wild type, TR_34_, and TR_46_, and *sdh*B alleles, split on H270Y allele, were analysed separately. We also analysed the fitness data from each of the three fungicides in separate models. The no-fungicide treatment was included in all of these models as concentration 0. We specified Malthusian fitness as the response variable and variant, fungicide concentration, and the interaction between the two as explanatory variables. Because the fitness values of different alleles from the same bottle are not independent, the sample barcode was included as a sample-level random effect. Model assumptions of homoscedasticity and normality of residuals were checked for all models. We calculated the marginal means of the treatment groups using the emmeans package.^42^ Among the grass samples, we tested for post hoc pairwise differences between the alleles in each fungicide treatment using *t*-tests on estimated marginal means with Bonferroni correction for multiple testing. The compact-letter displays used to indicate statistical significance between groups were generated based on Sidak-adjusted groupings. We only had agar samples at the highest fungicide concentrations. Therefore, when modelling the effect of agar versus grass as a substrate on the fitness of the alleles, we only included grass samples matching the fungicide concentrations of the agar samples for each of the three fungicides. We fitted our linear mixed-effects models with variant, growth substrate, and the interaction between the two as explanatory variables. Again, the sample barcode was included as a sample-level random effect. As a post hoc test, we examined pairwise differences between grass and agar samples for each variant using *t*-tests on estimated marginal means with Bonferroni correction for multiple testing. All data figures were generated using the ggplot2 package^43^ and cowplot packages.^44^ The packages from the tidyverse were used for data wrangling.

## Supplementary Figures and Tables

**Figure S1.**
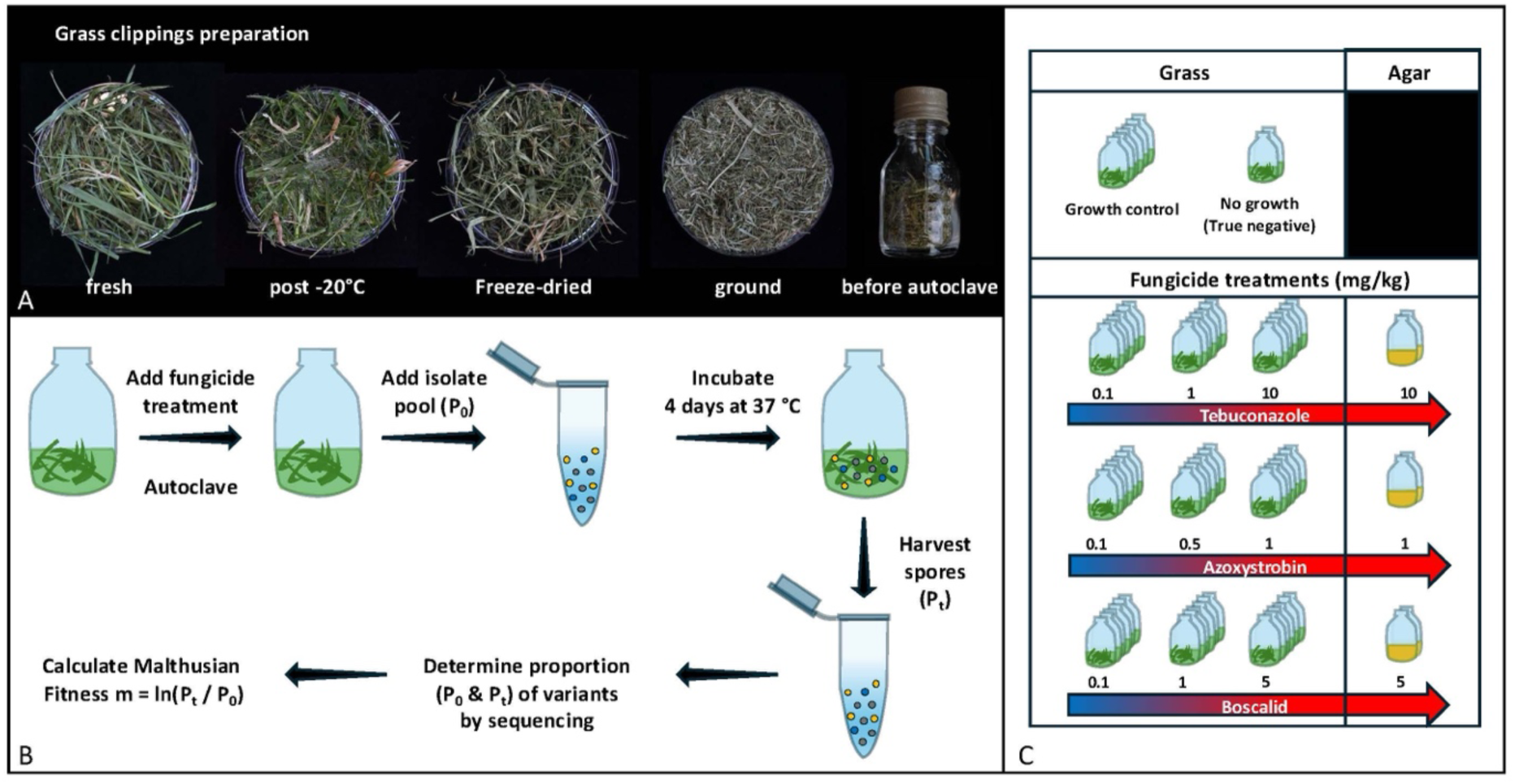
Schematic overview of experimental grass mesocosm design. A) The different steps to prepare to fresh grass clippings for the grass mesocosms experiments. Fresh fungicide free grass clippings are stored at -20 °C within 24 hours of collection. After thawing, the grass is freeze-dried and subsequently sieved and grinded through a 1 cm filter, 1 gr of ground grass is used in a 30mL glass bottle. B) Fungicide solutions are added to their respective grass bottles to rehydrate the grass and subsequently autoclaved. The pool of *A. fumigatus* isolates is then added to each bottle. After incubation the spores are harvested, and the proportion of variants is determined by sequencing. P_0_ and P_t_ are used to calculate the Malthusian fitness of the alleles in for samples. C) The fungicides and their concentrations used in this study for the grass and the agar bottles.

**Figure S2.**
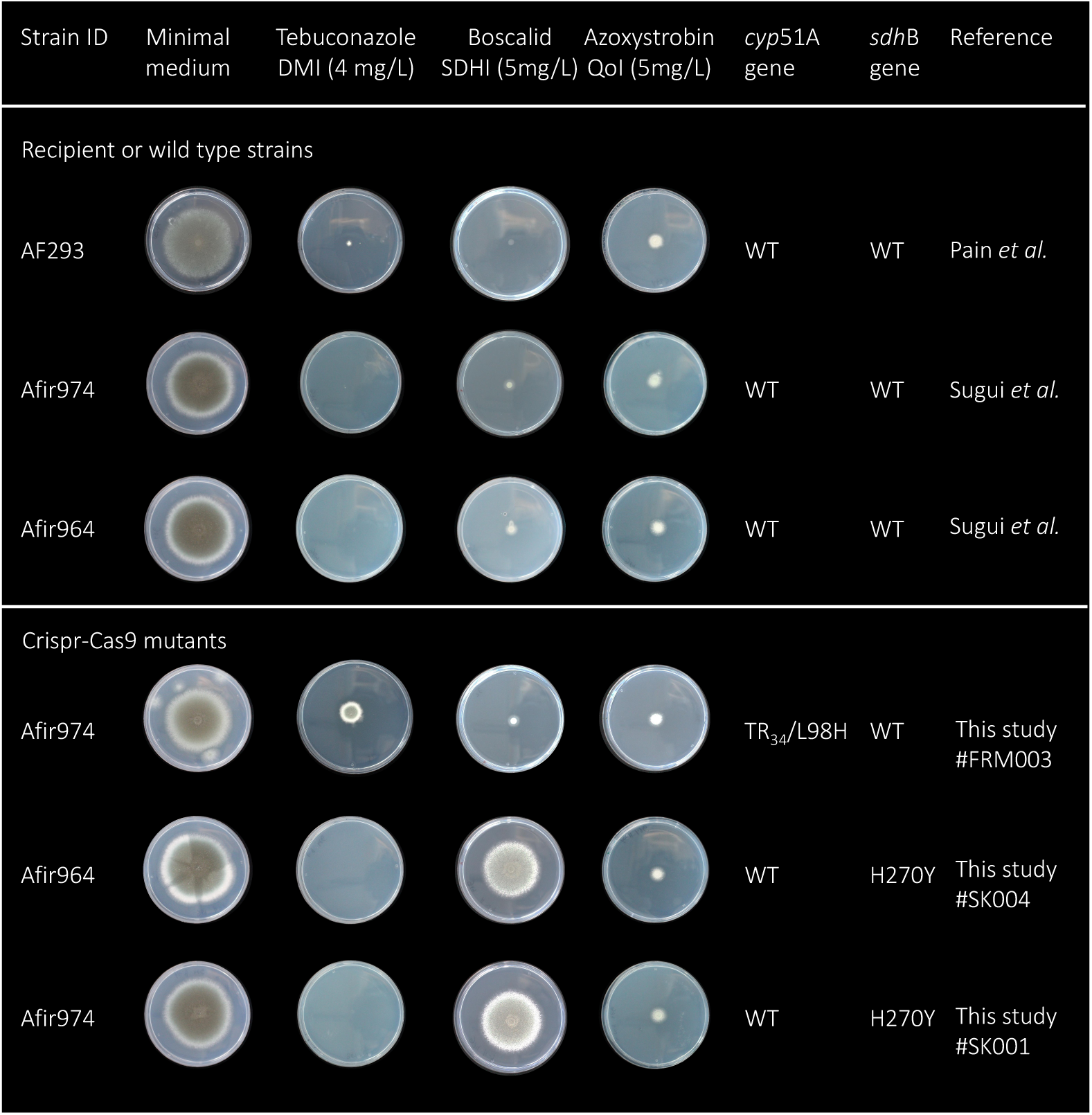
Single resistance allele mutants and their fungicide susceptibility profile. Wild type isolate AF293 (Pain *et al.* FGB 2004; 41: 443–453) and CrisprCas9 mutant recipient strains Afir974 and Afir964 (Sugui *et al.* mBio 2011; 2 (6) e00234-11) are shown in the top panel with from left to right complete medium (CM), CM with tebuconazole (DMI), boscalid (SDHI) and azoxystrobin (QoI). All wild type isolates show no growth under fungicide selection and are susceptible. The Crispr-Cas9 mutants on the bottom panel only show growth and thereby phenotypic resistance to tebuconazole (DMI) in case of a *cyp*51A mutation (TR_34_L98H) and to boscalid (SDHI) in case of a *sdh*B mutation (H270Y). QoI resistance is correlated to a mutation in the *cyt*B gene (G143A) in the mitochondrial genome for which at this time no methodological approaches are available to construct mutants, the TR_34_L98H *cyp*51A and the H270Y *sdh*B resistance alleles do not give rise to azoxystrobin resistance.

**Figure S3.**
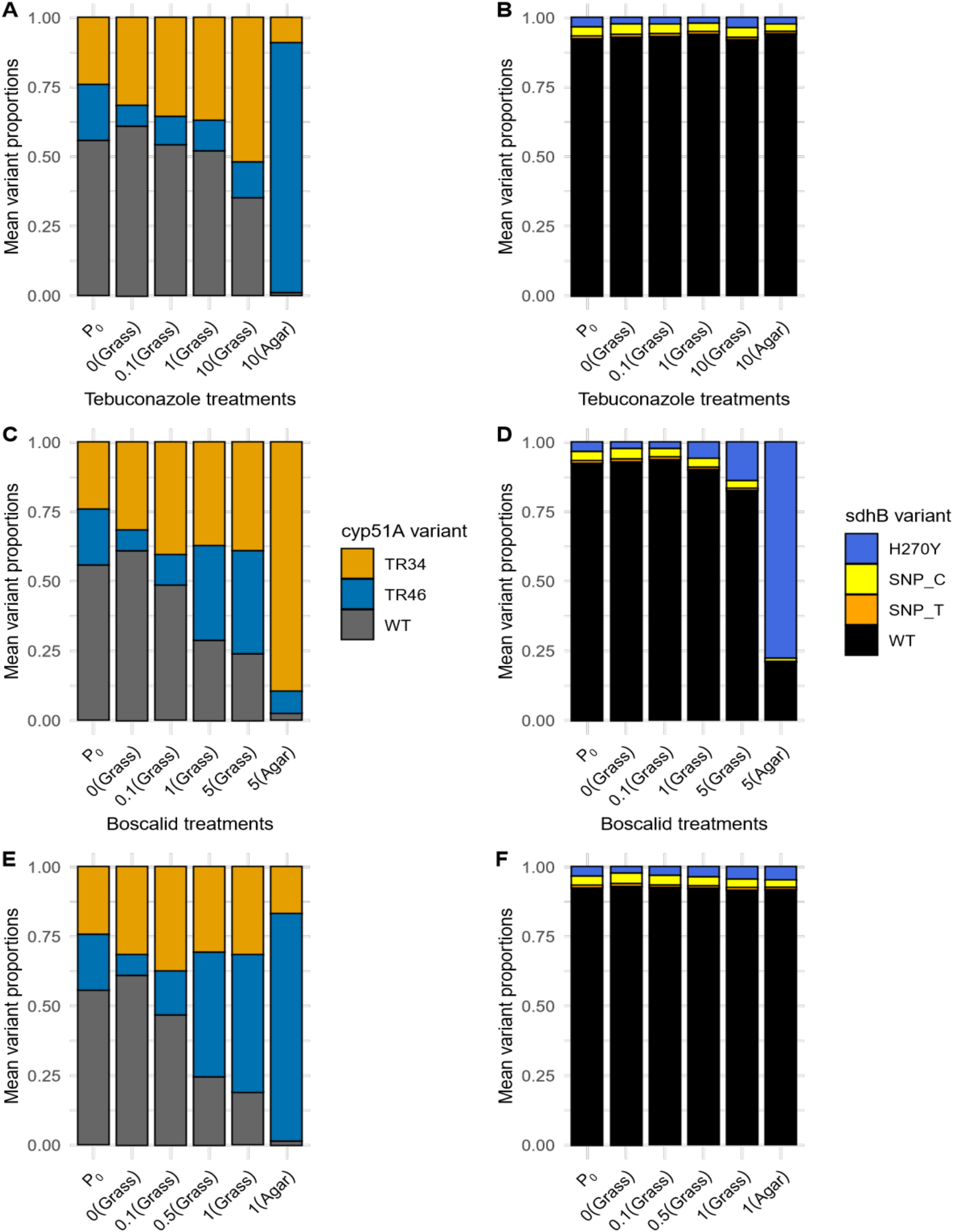
Variant proportions for *cyp*51A and *sdh*B. The proportions visualized in this figure were used to calculate the Malthusian fitness as shown in Figure 1 in the main manuscript.

**Figure S4.**
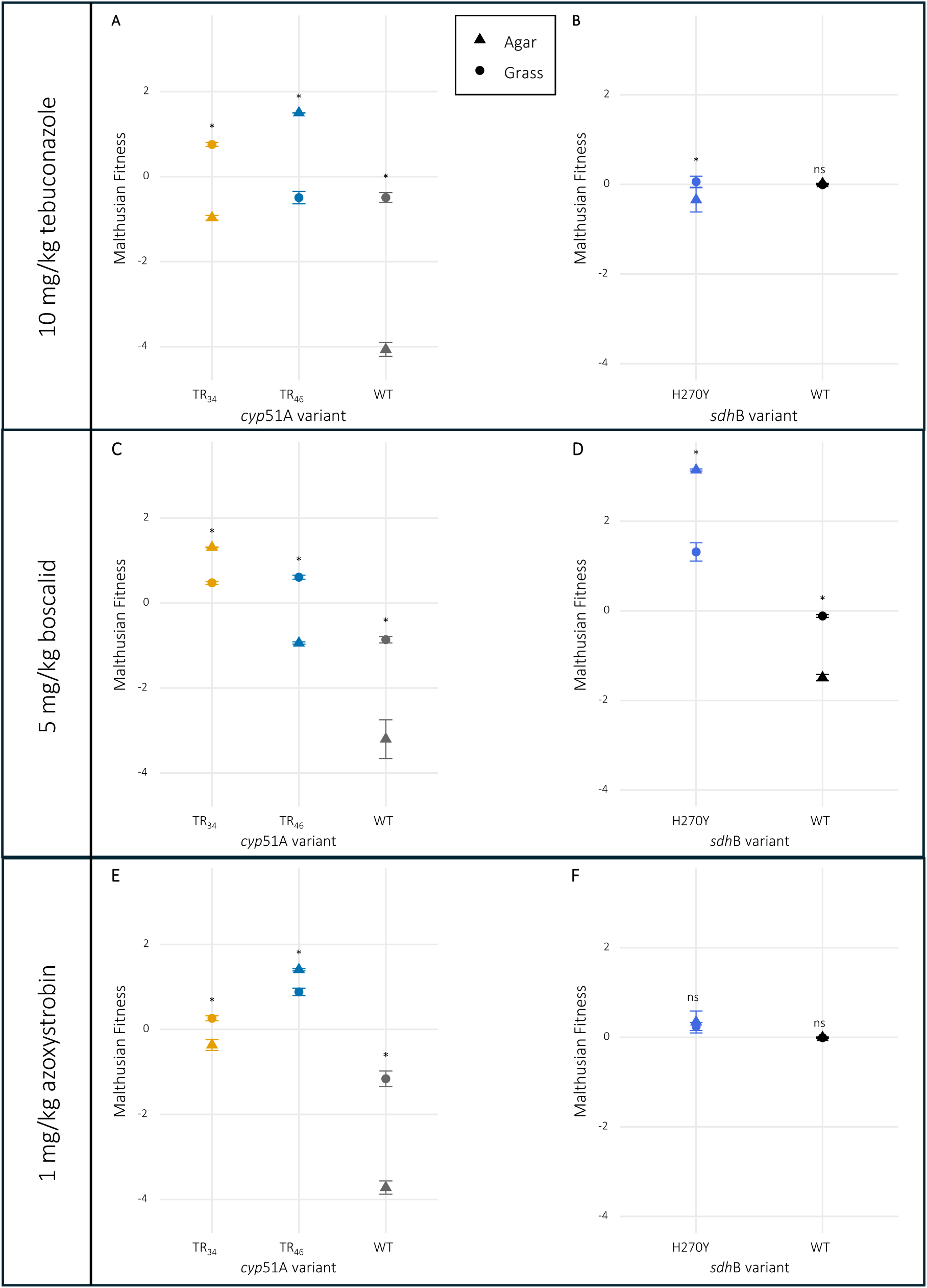
Fitness plot of fitness of the *cyp*51A and *sdh*B alleles at the highest fungicide concentrations comparing fitness on agar versus in the grass substrate per fungicide per variant. A) The fitness of the *cyp*51A alleles on 10 mg/kg tebuconazole. B) The fitness of the *sdh*B alleles on 10 mg/kg tebuconazole. C) The fitness of the *cyp*51A alleles on 5 mg/kg boscalid. D) The fitness of the *sdh*B alleles on 5 mg/kg boscalid. E) The fitness of the *cyp*51A alleles on 1 mg/kg azoxystrobin. F) The fitness of the *sdh*B alleles on 1 mg/kg azoxysytrobin. Shapes indicate the growth substrate. Post hoc pairwise comparisons were performed using *t*-tests on estimated marginal means with Bonferroni correction for multiple testing. (* = p < 0.05).

**Figure S5.**
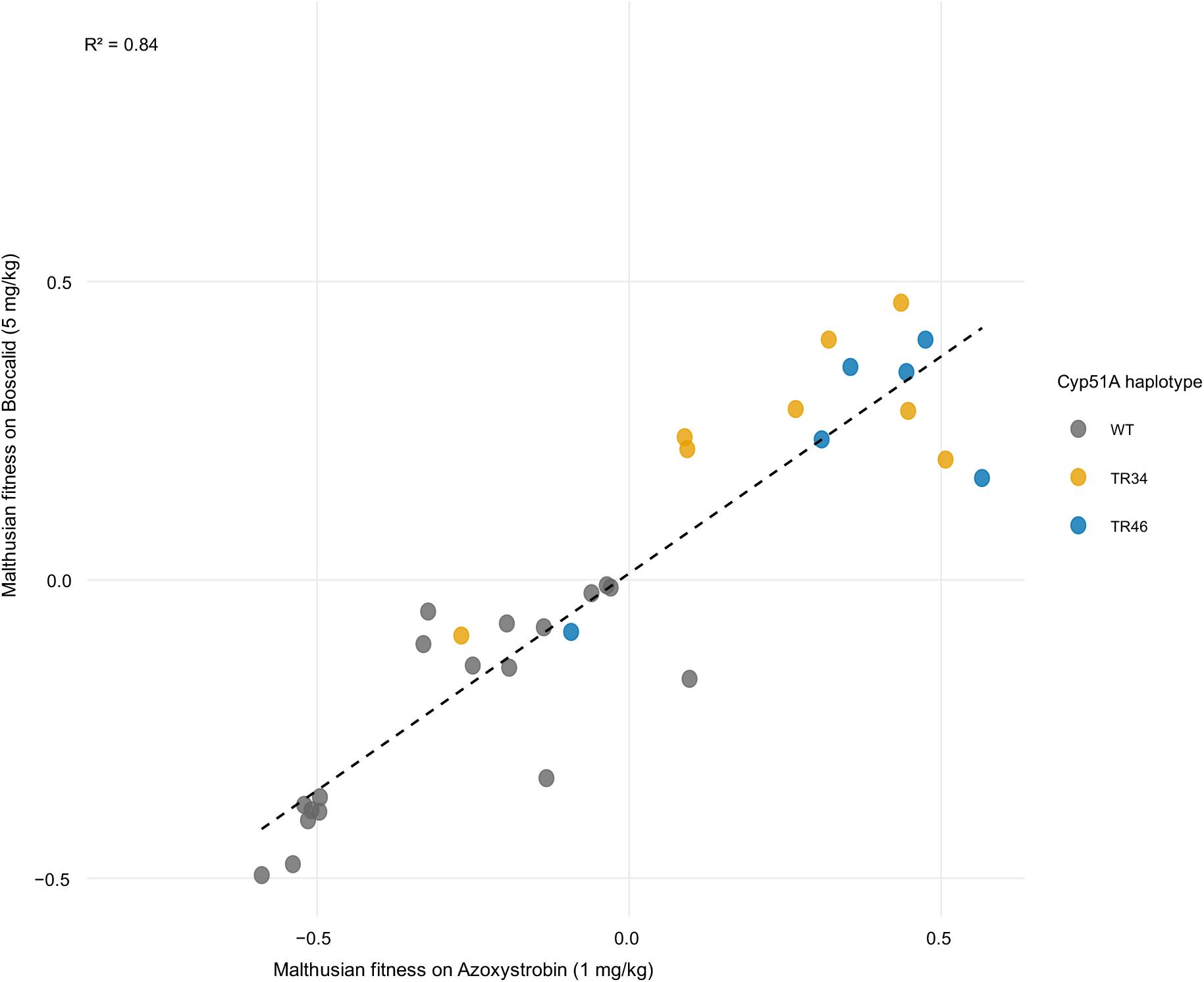
Correlation plot of the fitness on azoxystrobin (1 mg/kg) versus boscalid (5 mg/kg) of the *cyp*51A haplotypes, coloured by haplotype group. The fitness values correspond to those shown in Figure 2 in the main manuscript.

**Table S1.**
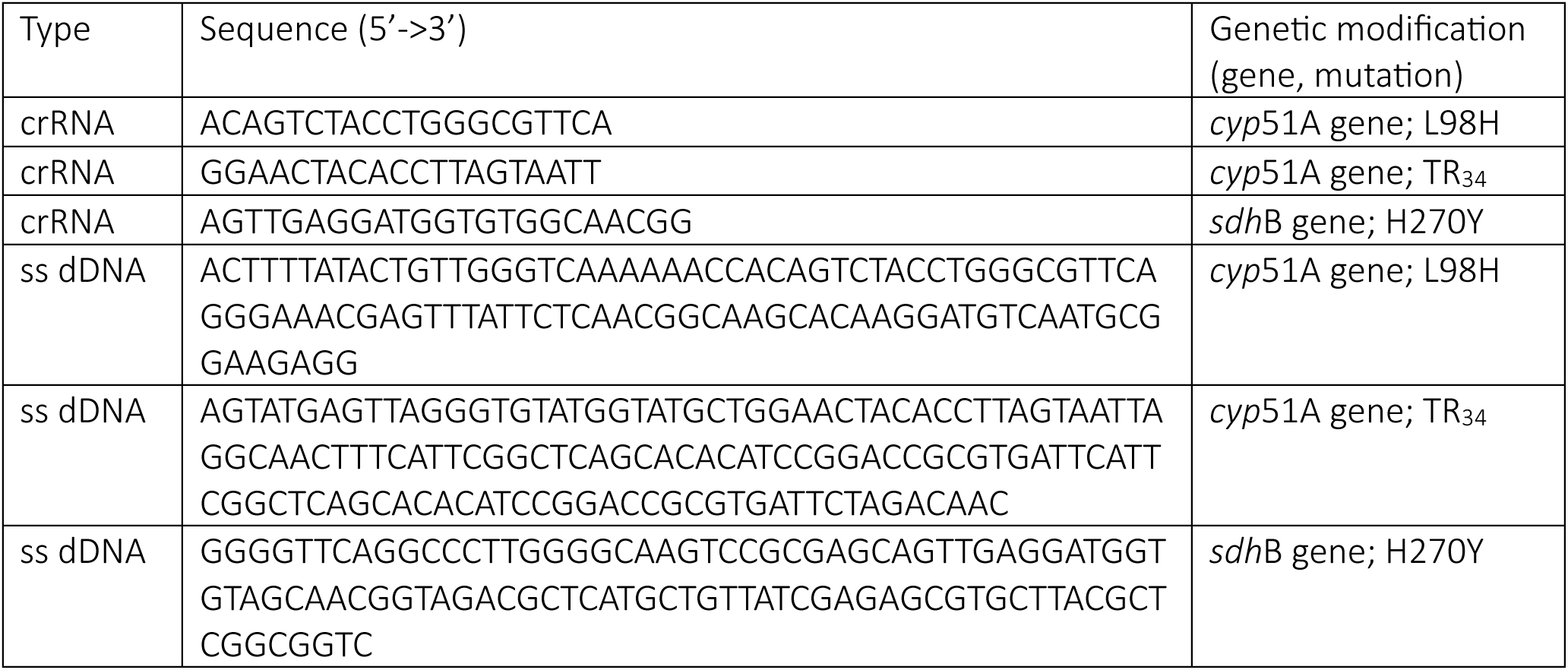
CRISPR cas9 mutant construction sequences. The following crisprRNA (crRNA) were used in combination with single stranded donor DNA (ss dDNA).

**Table S2.** Pairwise fitness differences between *cyp*51A alleles at different tebuconazole concentrations. This table matches the data of Figure 1A. Post hoc pairwise comparisons used *t*-tests on estimated marginal means with Bonferroni correction for multiple testing.

| Compound | Concentration<br>(mg/kg) | Contrast | Estimate | SE | df | t ratio | P value |
| --- | --- | --- | --- | --- | --- | --- | --- |
| Tebuconazole | 0.0 | TR<sub>34</sub>–TR<sub>46</sub> | 1.332 | 0.141 | 40 | 9.431 | < 0.001 |
|  |  | TR<sub>34</sub>–WT | 0.176 | 0.141 | 40 | 1.247 | 0.659 |
|  |  | TR<sub>46</sub>–WT | 1.156 | 0.141 | 40 | -8.184 | < 0.001 |
|  | 0.1 | TR<sub>34</sub>–TR<sub>46</sub> | 1.122 | 0.141 | 40 | 7.942 | < 0.001 |
|  |  | TR<sub>34</sub>–WT | 0.408 | 0.141 | 40 | 2.889 | 0.019 |
|  |  | TR<sub>46</sub>–WT | -0.714 | 0.141 | 40 | -5.053 | < 0.001 |
|  | 1.0 | TR<sub>34</sub>–TR<sub>46</sub> | 1.078 | 0.141 | 40 | 7.629 | < 0.001 |
|  |  | TR<sub>34</sub>–WT | 0.496 | 0.141 | 40 | 3.508 | 0.003 |
|  |  | TR<sub>46</sub>–WT | -0.582 | 0.141 | 40 | -4.121 | 0.001 |
|  | 10.0 | TR<sub>34</sub>–TR<sub>46</sub> | 1.250 | 0.141 | 40 | 8.852 | < 0.001 |
|  |  | TR<sub>34</sub>–WT | 1.249 | 0.141 | 40 | 8.842 | < 0.001 |
|  |  | TR<sub>46</sub>–WT | -0.001 | 0.141 | 40 | -0.010 | 1.000 |
| Azoxystrobin | 0.0 | TR<sub>34</sub>–TR<sub>46</sub> | 1.336 | 0.126 | 40 | 10.563 | < 0.001 |
|  |  | TR<sub>34</sub>–WT | 0.176 | 0.126 | 40 | 1.397 | 0.511 |
|  |  | TR<sub>46</sub>–WT | -1.156 | 0.126 | 40 | -9.167 | < 0.001 |
|  | 0.1 | TR<sub>34</sub>–TR<sub>46</sub> | 1.157 | 0.126 | 40 | 9.174 | < 0.001 |
|  |  | TR<sub>34</sub>–WT | 0.650 | 0.126 | 40 | 5.153 | < 0.001 |
|  |  | TR<sub>46</sub>–WT | -0.507 | 0.126 | 40 | -4.021 | 0.001 |
|  | 1.0 | TR<sub>34</sub>–TR<sub>46</sub> | -0.067 | 0.126 | 40 | -0.534 | 1.000 |
|  |  | TR<sub>34</sub>–WT | 1.145 | 0.126 | 40 | 9.077 | < 0.001 |
|  |  | TR<sub>46</sub>–WT | 1.212 | 0.126 | 40 | 9.611 | < 0.001 |
|  | 5.0 | TR<sub>34</sub>–TR<sub>46</sub> | -0.131 | 0.126 | 40 | -1.042 | 0.911 |
|  |  | TR<sub>34</sub>–WT | 1.339 | 0.126 | 40 | 10.613 | < 0.001 |
|  |  | TR<sub>46</sub>–WT | 1.470 | 0.126 | 40 | 11.655 | < 0.001 |
| Boscalid | 0.0 | TR<sub>34</sub>–TR<sub>46</sub> | 1.333 | 0.145 | 40 | 9.161 | < 0.001 |
|  |  | TR<sub>34</sub>–WT | 0.176 | 0.145 | 40 | 1.211 | 0.699 |
|  |  | TR<sub>46</sub>–WT | -1.156 | 0.145 | 40 | -7.950 | < 0.001 |
|  | 0.1 | TR<sub>34</sub>–TR<sub>46</sub> | 0.679 | 0.145 | 40 | 4.667 | < 0.001 |
|  |  | TR<sub>34</sub>–WT | 0.612 | 0.145 | 40 | 4.206 | < 0.001 |
|  |  | TR<sub>46</sub>–WT | -0.067 | 0.145 | 40 | -0.462 | 1.000 |
|  | 0.5 | TR<sub>34</sub>–TR<sub>46</sub> | -0.552 | 0.145 | 40 | -3.795 | 0.002 |
|  |  | TR<sub>34</sub>–WT | 1.115 | 0.145 | 40 | 7.663 | < 0.001 |
|  |  | TR<sub>46</sub>–WT | 1.667 | 0.145 | 40 | 11.458 | < 0.001 |
|  | 1.0 | TR<sub>34</sub>–TR<sub>46</sub> | -0.622 | 0.145 | 40 | -4.279 | < 0.001 |
|  |  | TR<sub>34</sub>–WT | 1.421 | 0.145 | 40 | 9.772 | < 0.001 |
|  |  | TR<sub>46</sub>–WT | 2.044 | 0.145 | 40 | 14.051 | < 0.001 |

WT: wild type

**Table S3.**
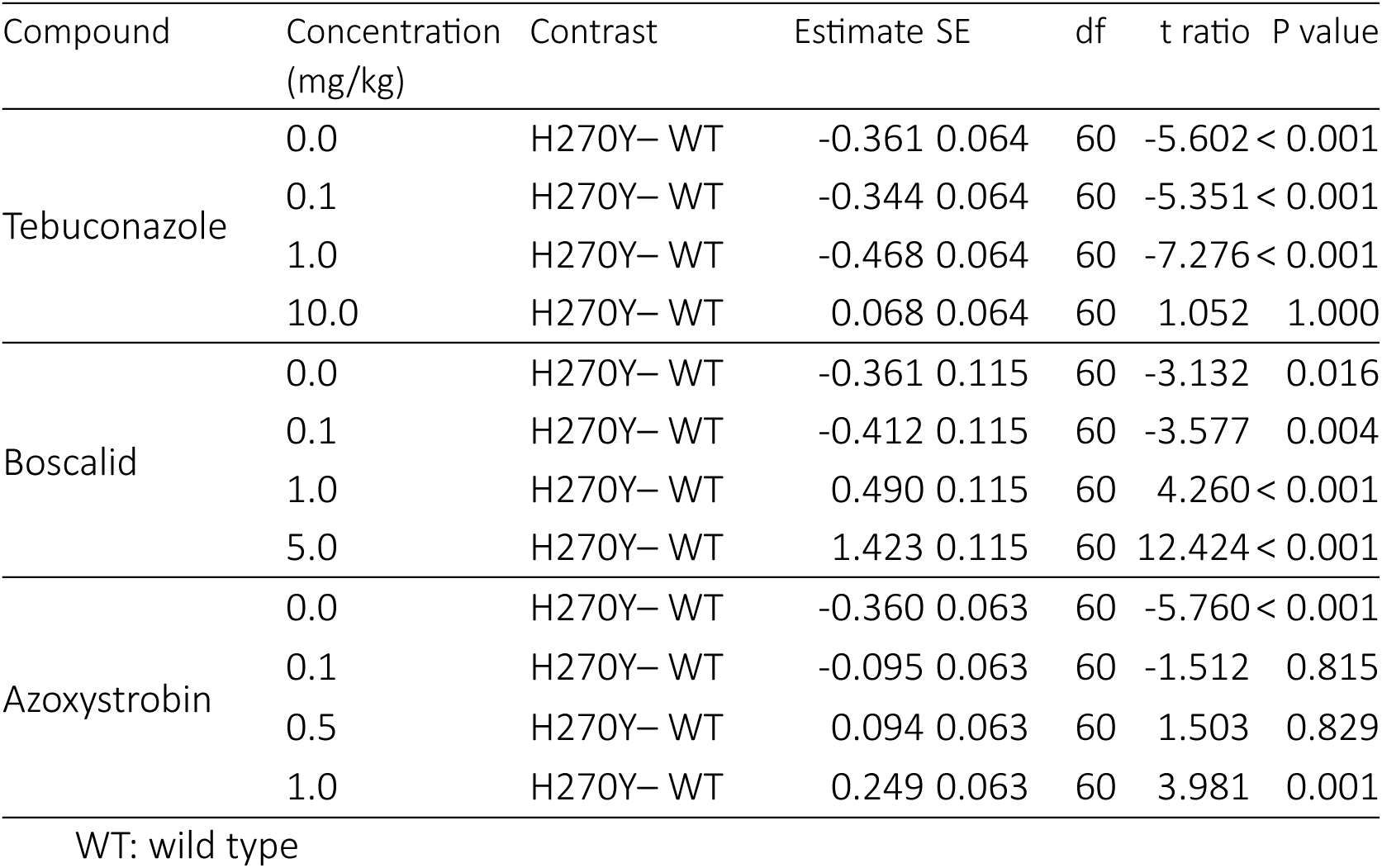
Pairwise fitness differences between *sdh*B alleles at different fungicide concentrations. This table matches the data of Figure 1B. Post hoc pairwise comparisons used *t*-tests on estimated marginal means with Bonferroni correction for multiple testing.

**Table S4.**
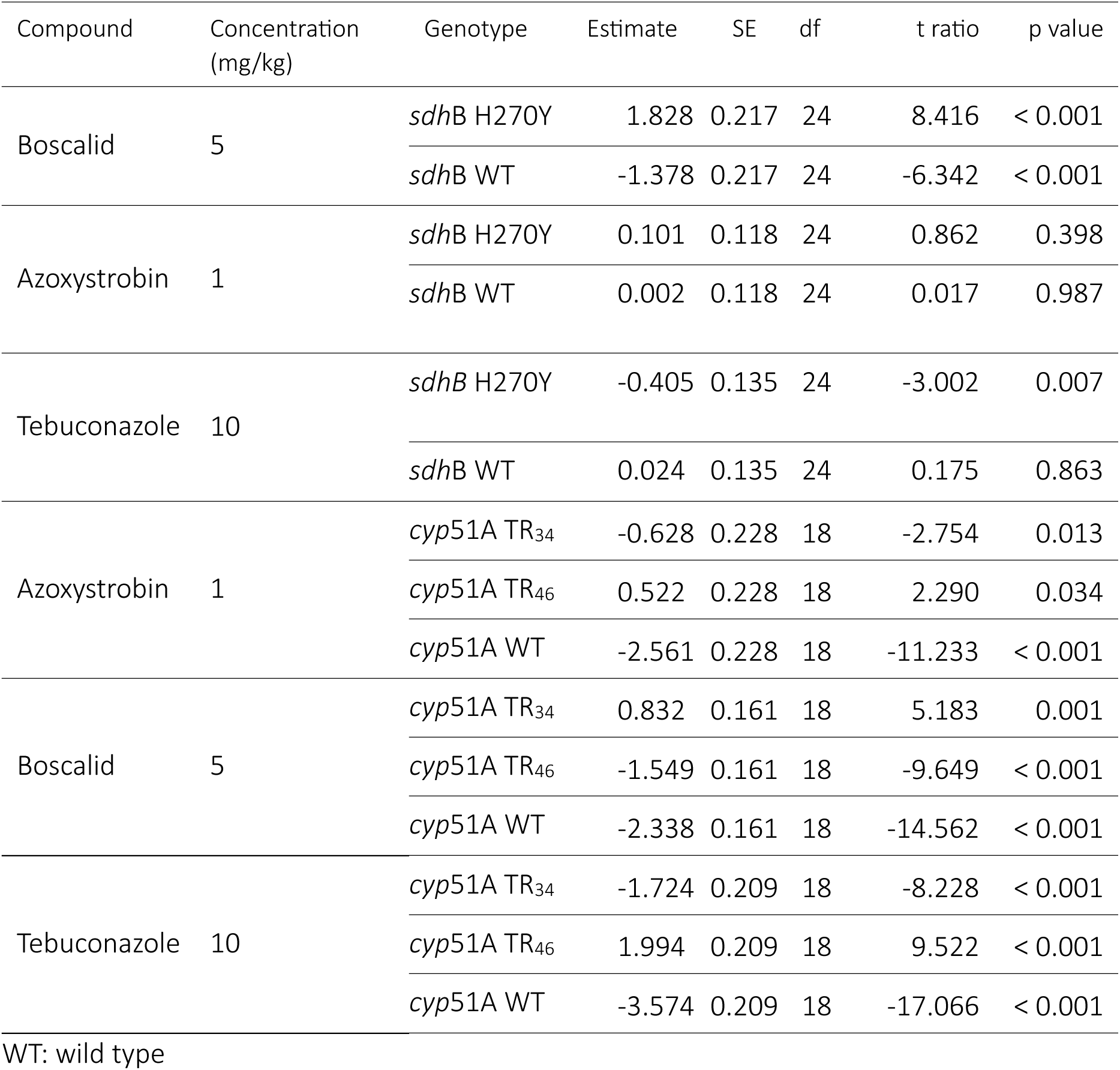
Pairwise differences in fitness between agar and grass competitions. Estimates represent the contrast Agar – Grass within each genotype. Post hoc pairwise comparisons were performed using *t*-tests on estimated marginal means with Bonferroni correction for multiple testing.

| Compound | Concentration<br>(mg/kg) | Genotype | Estimate | SE | df | t ratio | p value |
| --- | --- | --- | --- | --- | --- | --- | --- |
| Boscalid | 5 | *sdhB* H270Y | 1.828 | 0.217 | 24 | 8.416 | < 0.001 |
|  |  | *sdhB* WT | -1.378 | 0.217 | 24 | -6.342 | < 0.001 |
| Azoxystrobin | 1 | *sdhB* H270Y | 0.101 | 0.118 | 24 | 0.862 | 0.398 |
|  |  | *sdhB* WT | 0.002 | 0.118 | 24 | 0.017 | 0.987 |
| Tebuconazole | 10 | *sdhB* H270Y | -0.405 | 0.135 | 24 | -3.002 | 0.007 |
|  |  | *sdhB* WT | 0.024 | 0.135 | 24 | 0.175 | 0.863 |
| Azoxystrobin | 1 | *cyp51A* TR<sub>34</sub> | -0.628 | 0.228 | 18 | -2.754 | 0.013 |
|  |  | *cyp51A* TR<sub>46</sub> | 0.522 | 0.228 | 18 | 2.290 | 0.034 |
|  |  | *cyp51A* WT | -2.561 | 0.228 | 18 | -11.233 | < 0.001 |
| Boscalid | 5 | *cyp51A* TR<sub>34</sub> | 0.832 | 0.161 | 18 | 5.183 | 0.001 |
|  |  | *cyp51A* TR<sub>46</sub> | -1.549 | 0.161 | 18 | -9.649 | < 0.001 |
|  |  | *cyp51A* WT | -2.338 | 0.161 | 18 | -14.562 | < 0.001 |
| Tebuconazole | 10 | *cyp51A* TR<sub>34</sub> | -1.724 | 0.209 | 18 | -8.228 | < 0.001 |
|  |  | *cyp51A* TR<sub>46</sub> | 1.994 | 0.209 | 18 | 9.522 | < 0.001 |
|  |  | *cyp51A* WT | -3.574 | 0.209 | 18 | -17.066 | < 0.001 |

WT: wild type

